# Heart rate synchrony as a marker of real-world social engagement: modulated by proximity, social familiarity, and acoustic environment

**DOI:** 10.64898/2025.12.01.691366

**Authors:** Hanlu He, Jeppe Høy Christensen, A. Josefine Munch Sørensen, Ivana Konvalinka

**Author notes:** Contributing authors.

## Abstract

Human social behaviour unfolds in complex real-world environments influenced by social and environmental factors, yet reliable markers of social engagement and connection remain elusive. Interpersonal physiological synchrony (IPS) has been proposed as one such marker, but its occurrence in everyday settings is not well established. To investigate the social and environmental factors that influence IPS, we continuously measured heart rate, GPS, and the acoustic environment from 72 participants across three multi-day trips to New York City, capturing naturalistic social behaviour. Across all three trips, heart rates reliably synchronized when participants were in close physical proximity, indicating that shared environmental context was sufficient to elicit IPS. IPS was stronger among socially familiar peers, and context dependent, emerging during close-proximity interactions and joint attention to shared stimuli, but not dispersed interactions. IPS was also modulated by acoustic context, with low-to-moderate sound pressure levels and moderate-to-high signal-to-noise ratios enhancing synchrony, while excessive environmental noise reduced it to levels comparable to non-interactive settings, which may reflect lower levels of joint engagement in noisier environments. These findings demonstrate that IPS emerges in naturalistic social settings and is modulated by physical proximity, social familiarity, social context, and the acoustic environment, establishing IPS as a reliable marker of real-world social engagement.

## 1 Introduction

Human social behaviour unfolds in complex, dynamic real-world environments, where individuals continuously coordinate their actions, perceptions, and internal physiological signals during shared experiences. One well-documented form of such coordination is interpersonal synchrony, which spontaneously emerges between people’s movements and physiological signals when they interact with one another[1–4], when they passively perceive each other[5] or when they merely have the same sensory input[6–8]. For example, people unconsciously synchronize their strides when walking side-by-side[9], and audiences spontaneously synchronize their clapping following a theater or opera performance[10]. Similarly, interpersonal physiological synchrony (IPS) - simultaneous changes in heart rates or phase-coupling between people’s periodic cardiac rhythms - accompanies behavioural coordination during face-to-face interactions between mothers and infants and among romantic or marital partners[9, 11, 12], as well as during shared coordinated experiences such as choir singing[13] or playful joint actions[14, 15]. IPS can emerge even without overt behavioural coordination, as shown by the synchronization of heart rhythms between performers and socially affiliated spectators during collective rituals such as fire-walking[16].

Despite widespread evidence for interpersonal synchrony across bodily signals and social contexts, the mechanisms underlying IPS and its potential functional role remain elusive, particularly within the complex, multisensory environments in which everyday social interactions occur. Prior research has consistently linked behavioural synchrony to social bonding and prosocial behaviour, showing that people like each other more, feel greater affiliation, and act more cooperatively after synchronizing their movements[17–21]. Similarly, IPS has been shown to be enhanced by social affiliation[22, 23]. For example, during a fire-walking ritual, heart-rate coordination was observed only between spectators and performers with pre-existing social ties, but not between unrelated dyads[16]. Likewise, socially close pairs exhibited stronger heart-rate synchrony than less close or unaffiliated pairs during a high-intensity haunted house experience[23], and autonomic synchrony between individuals meeting on blind dates has been shown to predict subsequent mutual attraction[24].

IPS can also arise without direct social interaction or shared social context, such as when people independently watch the same films or listen to the same narratives[25, 26]. In such cases, neural and physiological alignment is driven primarily by shared sensory input, and is amplified by simultaneous attentional engagement. For instance, distracted participants exposed to narrative stimuli exhibit reduced inter-subject correlation of heart rates[26]. Co-presence, such as when individuals watch the same movies together, further amplifies IPS[5]. This is consistent with accounts proposing that IPS and neural synchrony may be markers of joint attention or collective engagement[27–31], which are enhanced by shared social context[32, 33].

However, whether and how IPS emerges during everyday social interactions outside of controlled laboratory environments remains unclear. Measuring engagement in naturalistic settings is particularly important yet challenging, as subtle disengagement is difficult to detect and may indicate sensory difficulty, especially in individuals with hearing loss or degraded sensory input who experience increased cognitive demands in complex environments[34–36]. In the absence of overt behavioural synchrony, it is further unclear whether IPS emerges in low-arousal real-world settings or is instead driven by simultaneous surges in heart rates during high arousal events. Establishing whether IPS emerges during spontaneous face-to-face social exchanges and natural episodes of joint attention, and whether it varies with social affiliation and acoustic properties of the environment, is therefore essential for evaluating its utility as a marker of real-world social engagement.

To address these questions, we conducted a study in ecologically valid, real-world settings by continuously recording heart rates from groups of students engaged in various social activities across three separate 4-day trips to New York (Trip 1: N = 23, Trip 2: N = 24, Trip 3: N = 25), organized as part of Oticon’s annual Audioexplorer events[37]. Across the three trips, continuous monitoring generated a large volume of multimodal data, with approximately 293, 367, and 385 total hours recorded per day for Trips 1–3, respectively — equivalent to 14–17 hours of physiological, spatial, and acoustic data per participant per day. We investigated effects of physical proximity, activity type, social affiliation, and environmental noise on IPS. We categorized interactions into three distinct activity types: events involving shared sensory stimuli (e.g., attending the same lecture), face-to-face interactions (e.g., group games), and dispersed group interactions where the participants were together but not necessarily engaged in close interactions with all members of the group (e.g., having dinner across numerous small tables). In addition, we tracked participants’ real-time GPS coordinates to quantify physical proximity, in order to investigate whether IPS emerges more broadly when people are together, as a result of the shared environment.

We further examined whether social affiliation modulates IPS by comparing heart rate synchrony between dyads with pre-existing social ties versus those without, while controlling for physical proximity. Given the cognitive demands of processing auditory stimuli in noisy environments - which are often associated with heightened cognitive load and reduced listening engagement[38–40] - we examined how such acoustic challenges encountered during various activities in New York City impacted IPS. Listening engagement may diminish when the perceived value of communication does not outweigh the effort required to understand the other person. To capture the acoustic context, we employed Oticon’s hearing aids attached to participants’ clothing, enabling real-time recording of environmental sound features. We then investigated whether elevated sound pressure levels (SPL) and lower signal-to-noise ratios (SNR) in the sound environment, which may contribute to disengagement, attenuate IPS.

Specifically, we hypothesized that significant levels of IPS, operationalized using measures of inter-subject correlation of heart rates (ISC-HR)[26], emerge when people are in close physical proximity, and that IPS is modulated by the type of activity people engage in. We expected IPS to be higher during face-to-face interactions involving reciprocal exchange of social signals, as well as during joint attention episodes to shared sensory stimuli, compared to dispersed interactions or physical separation. In addition, we hypothesized that social affiliation amplifies ISC-HR through heightened shared engagement, resulting in higher IPS between socially affiliated individuals. Finally, we hypothesized that ISC-HR would be higher in auditory environments characterized by higher SNR and lower SPL, facilitating listening and social engagement. Thus, taking into account the physical and social proximity, the type of social activity and hence amount of shared sensory input, and the acoustic environment, we aimed to establish IPS more broadly as a socially and acoustically modulated marker of engagement in real-world environments.

## 2 Results

Across three separate trips to New York, groups of 23, 24, and 25 student participants, respectively, took part in scheduled activities and had free time to explore the city. Throughout each trip, participants wore hearing aids with microphones and Garmin wristbands synchronized to their mobile phones, which continuously measured features of the sound environment and heart rates at various locations (Figure 1). Data were collected continuously across four days for each trip. First, we examined whether inter-subject correlation of heart rates (ISC-HR), a proxy for IPS, was modulated by physical proximity between participants. GPS data were used to objectively determine the physical distance between individuals, and to assess ISC-HR when people were together in groups and in pairs. We further explored whether IPS was also modulated by social proximity, evaluated as pairs who knew each other beforehand and came to the trip in groups, while controlling for physical proximity. Finally, we investigated IPS across different planned activities to see whether IPS depended on the amount of shared social or environmental signals, such as settings with shared sensory information, close proximity face-to-face interactions, and dispersed interactions.

**Fig. 1:**
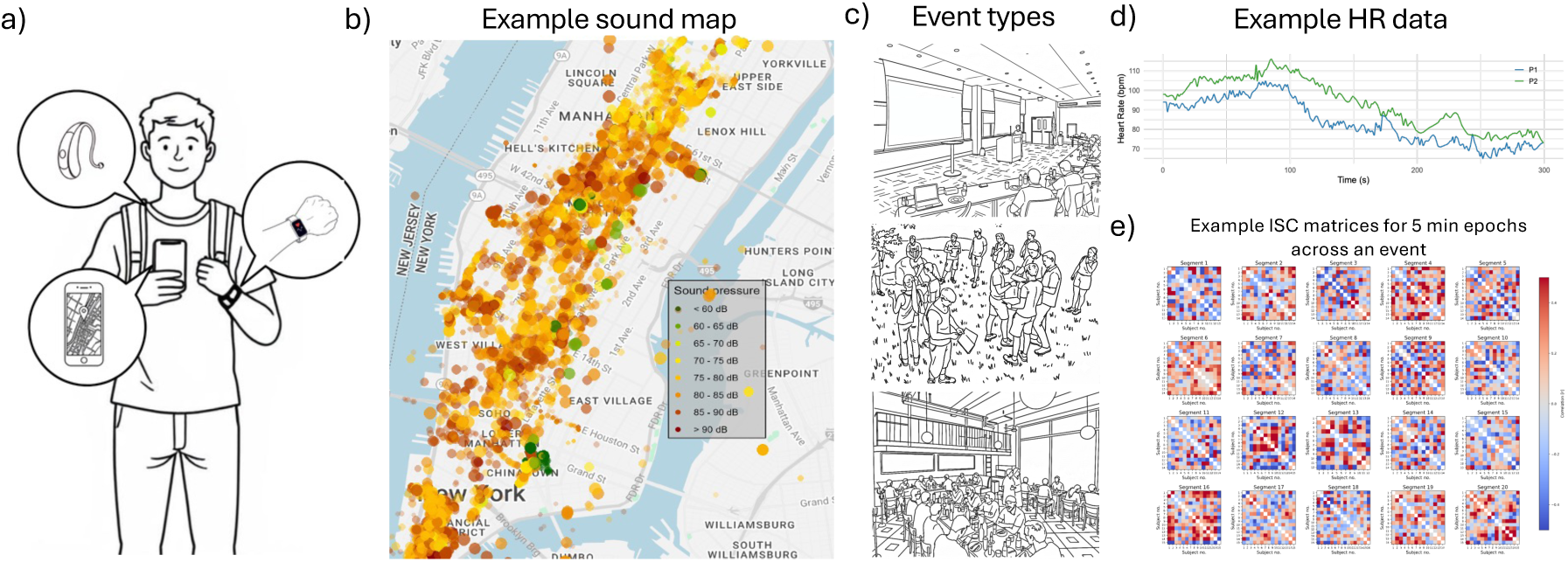
Study design and multimodal data collection during real-world interactions across three trips to New York City. **(a)** Participants wore wearable sensors throughout four-day trips to New York City: Garmin wristbands to measure heart rate (HR), hearing aid microphones clipped to the collar to estimate the sound environment, and mobile phones to record GPS coordinates and synchronize sensor streams. **(b)** Example of a “sound-stress” map generated from hearing aid microphones, with colored dots indicating sound pressure levels (SPL) across locations in the city. **(c)** Example illustrations of the three main event types: a lecture with shared sensory input (stimulus-locked), outdoor group work in Central Park (close proximity face-to-face interaction), and a restaurant setting (dispersed social interaction). **(d)** Example of continuous HR time series from two participants over a 5-minute window. **(e)** Inter-subject correlation of heart rate (ISC-HR) was computed by correlating HR time series across participants in 5-minute epochs, illustrated here as correlation matrices across an event. Across three separate trips (n = 23, 24, and 25 participants), data were collected continuously from 9:00–22:00 each day. This enabled us to examine how interpersonal physiological synchrony (IPS), indexed by ISC-HR, varied as a function of shared sensory environments, face-to-face proximity, and more dispersed interactions, while controlling for the acoustic environment and physical proximity of participants.

### 2.1 People’s heart rates synchronize when they are together in close physical proximity

To examine whether ISC-HR arises from mere physical co-presence, independent of social contexts [5], we used GPS data to identify periods when participants were physically close (within 20 m of each other) during daily activities (9:00–22:00) and segmented them into 5-minute windows. Across all three trips, ISC-HR was significantly higher during close-proximity group periods compared to time-misaligned shuffled controls, reflecting IPS between time-misaligned signals, with large effect sizes (Trip 1: *t*(22) = 14.36, *p* = 1.18 × 10*^−^*^1^^2^, *d* = 2.94; Trip 2: *t*(23) = 9.29, *p* = 2.98 × 10*^−^*^9^, *d* = 1.9; Trip 3: *t*(24) = 6.76, *p* = 5.41 × 10*^−^*^7^, *d* = 1.35) (Figure 2a)).

**Fig. 2:**
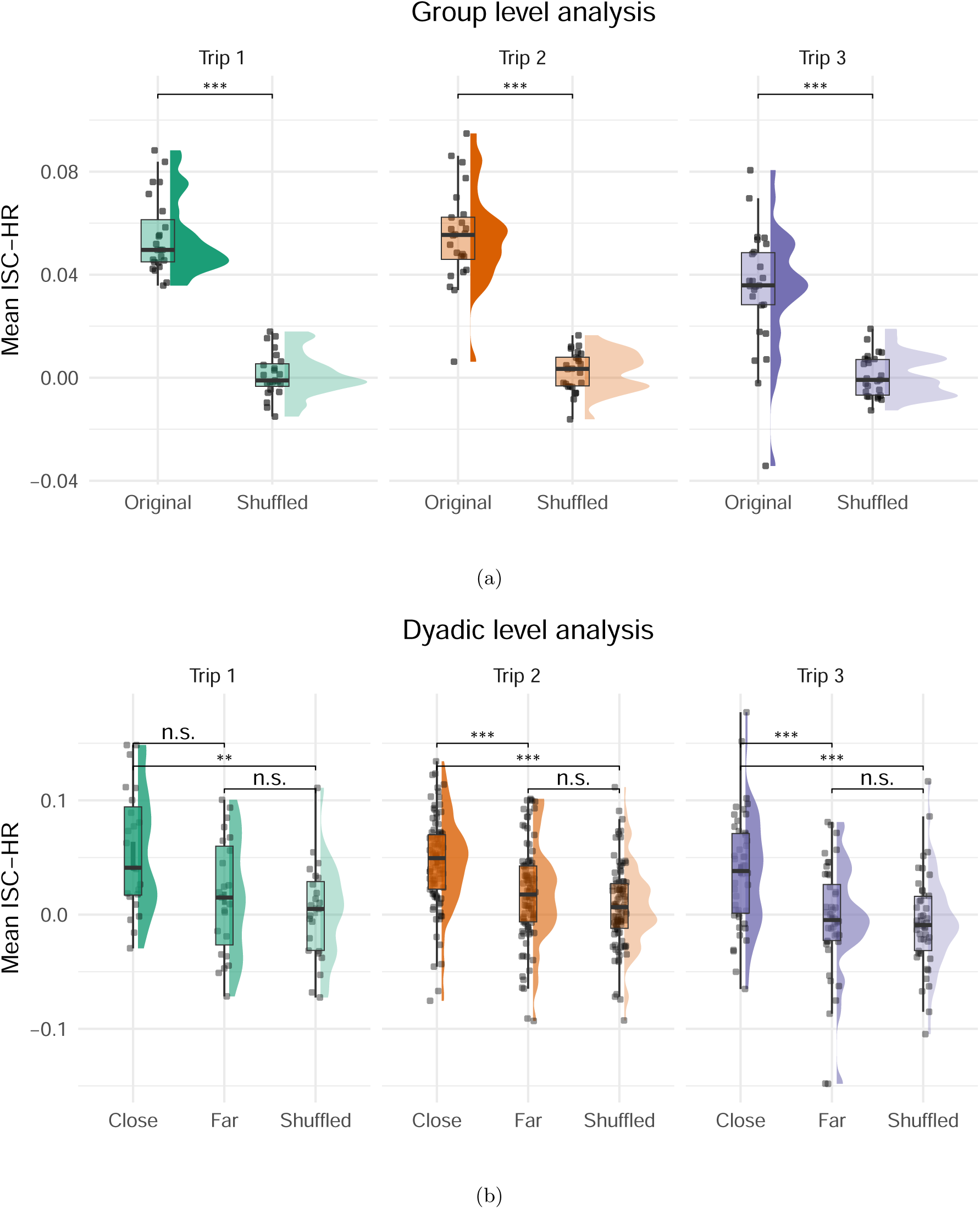
Heart rate synchrony emerges when people are in close physical proximity. **(a)** Group-level ISC-HR across all 5-minute segments when participants were physically together, defined using GPS proximity data during daily activities (9:00–22:00). Average ISC-HR values were significantly greater than those obtained from time-misaligned shuffled controls (see Section 4.4) across all three trips. **(b)** Pairwise ISC-HR comparisons under three physical proximity conditions: close (*<*20 m), far apart (*>*1 km), and time-misaligned shuffled control. Across all three trips, ISC-HR was significantly higher when individuals were in close proximity compared to time-misaligned shuffled controls, with moderate to large effect sizes. In Trips 2 and 3, ISC-HR was also significantly higher in close proximity compared to far proximity. No differences were observed between far and time-misaligned shuffled conditions across all trips. These findings show that IPS emerges between people in close physical proximity, both at the group and dyadic level. Significant differences are denoted with asterisks (∗ : *p <* 0.05, ∗∗ : *p <* 0.01, ∗ ∗ ∗ : *p <* 0.001).

We then examined ISC-HR at the dyadic level to better control for the physical proximity. Each pair was compared across three conditions: close proximity (within 20 m), far apart (more than 1 km), and time-misaligned shuffled controls. Repeated-measures ANOVAs revealed a consistent main effect of proximity across all trips (Trip 1: *F* (2, 44) = 7.21, *p* = 0.002, partial *η*^2^ = 0.18; Trip 2, *F* (2, 184) = 20.48, *p* = 9.29 × 10*^−^*^9^, partial *η*^2^ = 0.13; Trip 3, *F* (2, 88) = 8.92, *p* = 0.001, partial *η*^2^ = 0.12.). Post hoc tests confirmed that ISC-HR was reliably higher when pairs were in close proximity compared to time-misaligned shuffled controls (Trip 1: *t* = 4.03, *p*_Bonf._ = 0.002, *d* = 1.08; Trip 2: *t* = 6.26, *p*_Bonf._ = 3.54 × 10*^−^*^8^, *d* = 1; Trip 3: *t* = 3.51, *p*_Bonf._ = 0.003, *d* = 0.74).

For comparisons between close and far conditions, Trips 2 and 3 showed significant differences (Trip 2: *t* = 4.24, *p*_Bonf._ = 1.61 × 10*^−^*^4^, *d* = 0.57; Trip 3: *t* = 3.4, *p*_Bonf._ = 0.004, *d* = 0.79), while for Trip 1, the difference was not significant (*t* = 2.32, *p*_Bonf._ = 0.09, *d* = 0.76).Further, no significant differences emerged between the far and time-misaligned shuffled conditions (Trip 1: *t* = −1.18, *p*_Bonf._ = 0.75, *d* = −0.34; Trip 2: *t* = −2.3, *p*_Bonf._ = 0.07, *d* = −0.35; Trip 3: *t* = 1.55, *p*_Bonf._ = 0.38, *d* = 0.3) (Figure 2b).

Together, these results demonstrate that heart rate synchrony reliably emerges when individuals are in close physical proximity, both at the group and dyadic level, suggesting that shared environmental input facilitates physiological synchrony in real-world environments.

### 2.2 ISC-HR is modulated by social familiarity

The effect of social familiarity — defined as whether participants knew each other prior to the trip — was examined to test whether ISC-HR was modulated by pre-existing relationships beyond physical co-presence. Pairs were categorized as familiar (participants who knew and worked with each other before the trip) or unfamiliar (participants who were strangers prior to the trip). To control for physical proximity, pairwise ISC-HR values were computed only for segments when both participants were physically together identified by GPS coordinates in the same group and then averaged across those group segments.

The results revealed significantly higher ISC-HR for familiar pairs compared to unfamiliar pairs across all three trips (Trip 1: *W* = 3.45, *p* = 0.0029, rank-biserial *r* = 0.95, Trip 2: *W* = 4.68, *p* = 2.84 × 10*^−^*^6^, *r* = 0.32, Trip 3: *W* = 2.38, *p* = 0.018, *r* = 0.17), as seen in Figure 3. These findings indicate that social familiarity enhances interpersonal heart rate synchrony even when controlling for physical proximity, suggesting that pre-existing relationships facilitate physiological alignment.

**Fig. 3:**
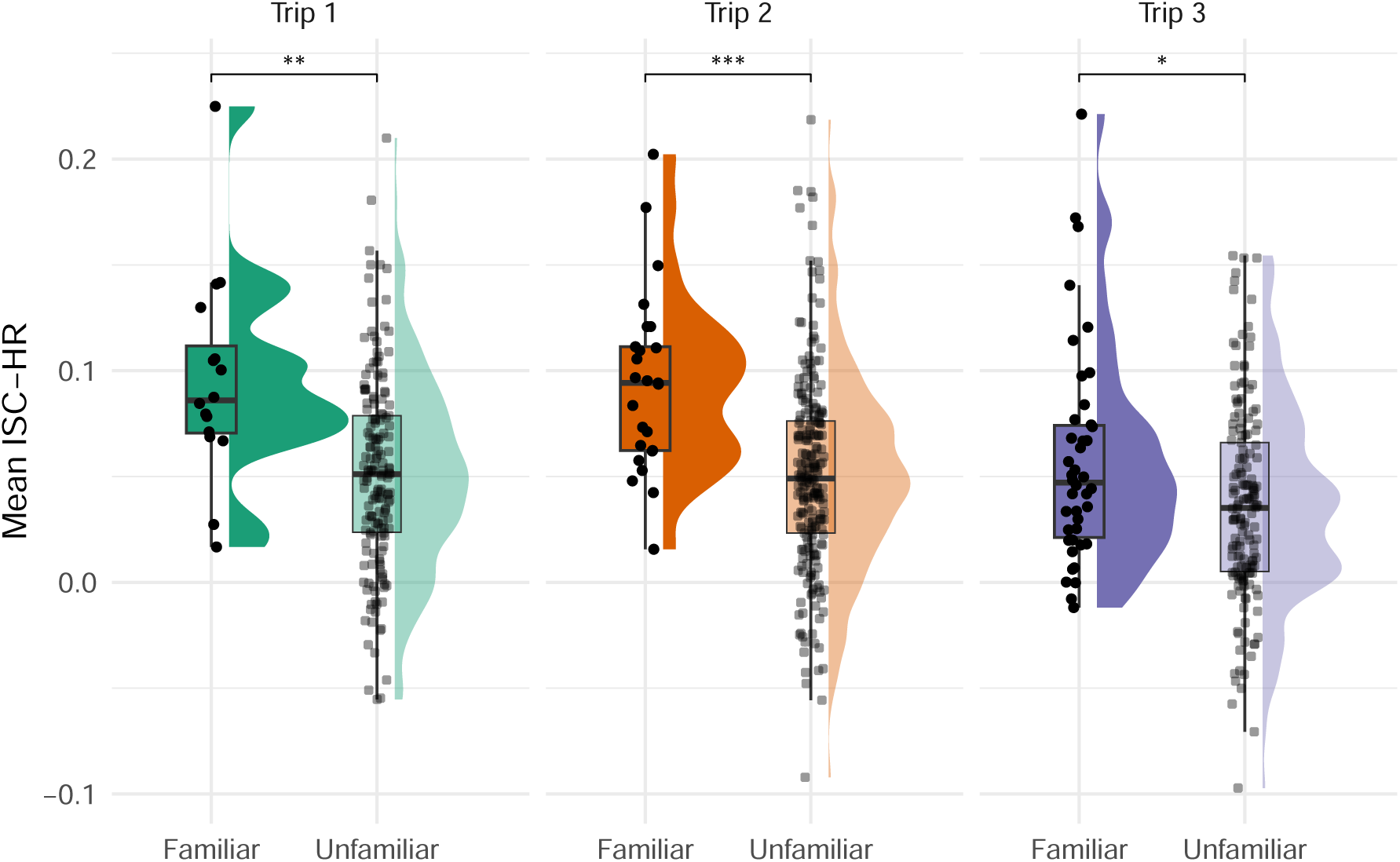
Comparison of ISC-HR between familiar and unfamiliar pairs across three trips. Mean ISC-HR values are shown for familiar pairs (participants who knew each other prior to the trip) and unfamiliar pairs (participants with no prior relationship), averaged across all group settings. Across all three trips, familiar pairs exhibited significantly higher ISC-HR than unfamiliar pairs, indicating that social proximity—defined by pre-existing social familiarity—enhances physiological synchrony beyond the effect of physical co-presence. Significant differences are denoted with asterisks (∗ : *p <* 0.05, ∗∗ : *p <* 0.01, ∗ ∗ ∗ : *p <* 0.001).

### 2.3 ISC-HR across different social contexts

To examine how different types of interactions relate to ISC-HR, events were first manually labeled based on their social and contextual characteristics. Because activities varied widely in structure and social engagement, they were grouped into three categories reflecting shared behavioral and attentional contexts: close-proximity interactions (e.g., group discussions during games or meals), where participants actively engaged in face-to-face interactions; stimulus-locked interactions (e.g., attending lecture presentations or shows), where everyone jointly attended to an external sensory input; and dispersed interactions (e.g., receptions or co-located activities without shared engagement), where participants were more scattered and interactions were brief (Figure 4).

**Fig. 4:**
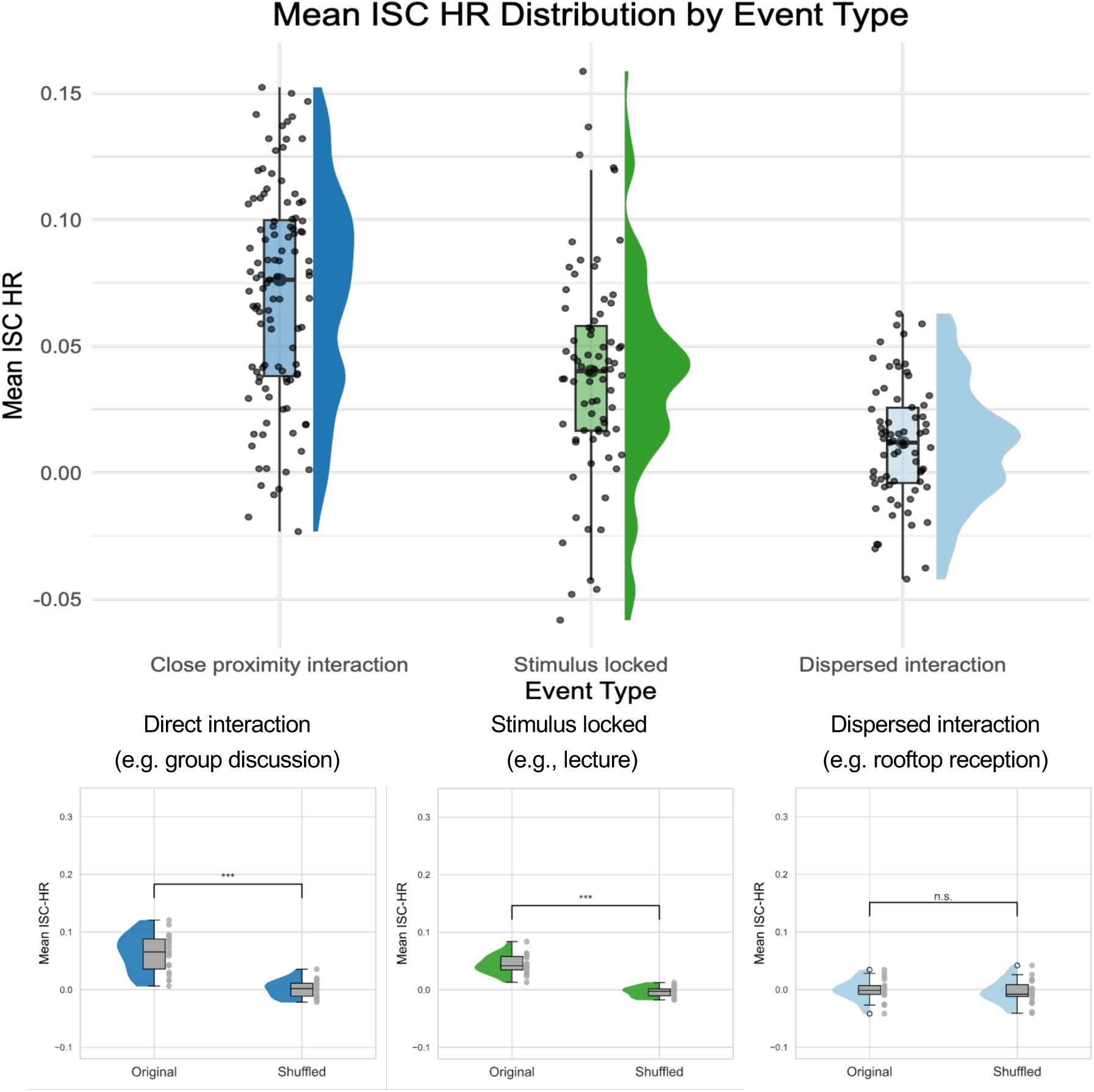
ISC-HR across different social interaction contexts. (**A**) Distribution of mean ISC-HR across participants during three types of interaction: close proximity interaction (e.g., group discussion or games), stimulus-locked interaction (e.g., attending a performance or lecture), and dispersed interaction (e.g., rooftop reception). ISC-HR was highest during close proximity interactions, intermediate during stimulus-locked interactions, and lowest during dispersed interactions. (**B**) Example events corresponding to the three categories. Each dot represents the mean ISC-HR of one participant with all others across 5-minute segments within the event, compared against a time-misaligned shuffled control generated by random pairing of time segments. Significant ISC-HR was observed during close proximity and stimulus-locked events, but not during dispersed interactions. Asterisks denote significant differences (Wilcoxon signed-rank test with Bonferroni correction; ∗ : *p <* 0.05, ∗∗ : *p <* 0.01, ∗ ∗ ∗ : *p <* 0.001).

Across all three trips, significant group-level ISC-HR was observed primarily during close-proximity and stimulus-locked events, but not during dispersed interactions. For example, in Trip 1, ISC-HR was significantly elevated compared to time-misaligned control during *Social Dinner 2*, where participants sat close together and engaged in conversation (*t* = 4.8, *p* = 2.9 × 10*^−^*^4^, *d* = 1.24), and during *Presentation 1*, where participants jointly attended a science lecture (*W* = 9.2, *p* = 1.26×10*^−^*^8^, *d* = 2). In contrast, no significant ISC-HR was observed for *Casual Reception 1*, where participants were scattered around a rooftop terrace (*t* = 1.09, *p* = 0.31, *d* = 0.36).

A similar pattern was found in Trip 2 and Trip 3, with presentation-based and interactive events in close proximity consistently eliciting high ISC-HR. For instance, in Trip 3, *Presentation 7* (joint attention to a shared stimulus) showed very strong HR synchrony (*t* = 11, *p* = 1.02 × 10*^−^*^8^, *d* = 2.33), as did *Group Game 2*, where participants engaged in intense group discussions to solve an engineering task (*t* = 5.51, *p* = 1.7 × 10*^−^*^4^, *d* = 1.26). In contrast, dispersed settings such as *Casual Dining 1–4* and *Casual Reception 1–2*, where participants were seated apart and loosely engaged, exhibited low and nonsignificant ISC-HR levels.

Importantly, these effects were not explained by differences in mean heart rate: across events, mean HR did not correlate with ISC-HR, indicating that synchrony reflects shared temporal fluctuations rather than global arousal levels (see S7 for control analyses). Full statistical details for all events are provided in the Supplementary Information (Figures S3–S5, Tables S2–S4).

### 2.4 The effect of Sound environment on ISC-HR

We examined whether exposure to different sound environments, characterized by sound pressure levels (SPL) and signal-to-noise ratios (SNR), was associated with variations in physiological synchrony. To contextualize the variability in acoustic sensory experience across the study, we visualized SPL and SNR over time for each trip. Figure 5a shows moment-to-moment and day-to-day fluctuations in the sound environment participants naturally encountered, with colour-coded background segments indicating when different social context event types occurred. Moreover, it shows that the sound environments were relatively consistent across the three trips.

**Fig. 5:**
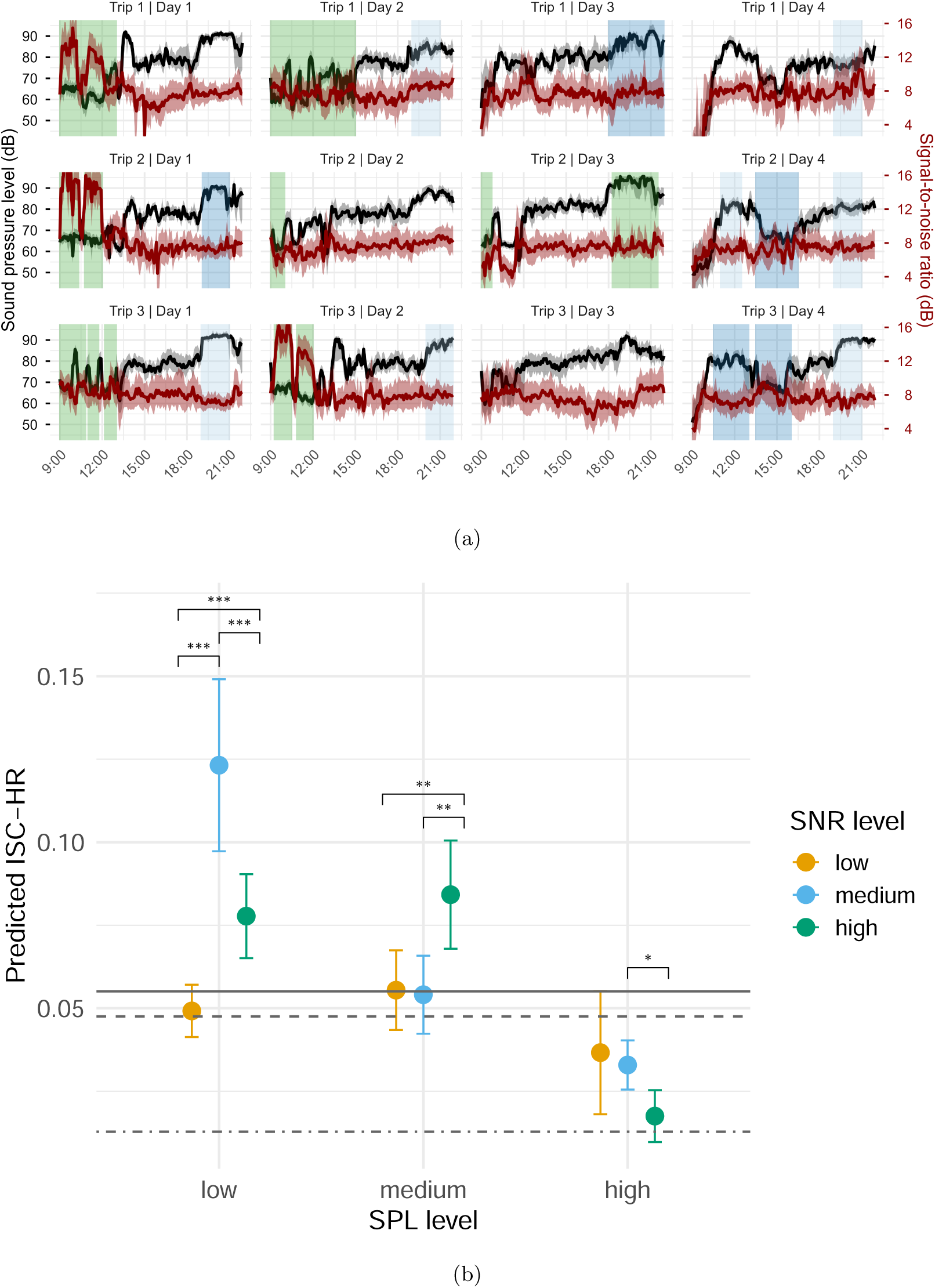
Sound environments and their relationship to ISC-HR. a) Median sound pressure level (left y-axis) and signal-to-noise ratio (right y-axis) in decibels for each trip (rows) and day (columns) across all participants estimated in 5-min windows. The shaded area around each value represents the interquartile range. Vertical shaded areas represent periods of different social interaction contexts (close proximity interaction = blue, stimulus locked = green, dispersed interaction = light blue). b) Predicted ISC-HR values across combinations of sound pressure level (SPL: low, medium, high) and signal-to-noise ratio (SNR: low, medium, high) estimated from a linear model including SPL, SNR, their interaction, and mean HR as a covariate. Points and error bars show estimated marginal means ± 95% CI for each SPL × SNR cell. Grey horizontal lines indicate mean ISC-HR observed in different event contexts (solid = close-proximity interaction, dashed = stimulus-locked, dot-dash = dispersed). Synchrony varied as a function of SPL and SNR: in quiet environments (low SPL), ISC-HR peaked at medium SNR; at moderate SPL, ISC-HR was strongest at high SNR, with no difference between low and medium SNR; in loud environments (high SPL), ISC-HR was generally lower, comparable to levels observed during dispersed interactions.

SPL and SNR values were estimated using participants’ hearing aids and categorized into low, medium, and high levels based on tertiles within participants. ISC-HR was modeled using a linear model including SPL level, SNR level, their interaction, and mean heart rate (HR) as a covariate (mean ISC ∼ SPL level × SNR level + mean HR). Inclusion of mean HR slightly improved explained variance (*R*^2^ increased from 0.0265 to 0.0286), but the effect was small, indicating that general arousal contributed minimally to ISC-HR.

Examination of the model revealed a significant interaction between SPL and SNR, indicating that the relationship between acoustic clarity and synchrony depends on the overall sound level. The following results describe this interaction in detail (Figure 5b).

#### Low SPL (quiet environments)

ISC-HR peaked at medium SNR, followed by high SNR, and was lowest for low SNR. Pairwise contrasts showed that medium SNR was significantly higher than low SNR (Δ^^^ = −0.0697, *SE* = 0.0106, *t*(5886) = −6.59, *p <* 0.0001) as well as high SNR (Δ^^^ = −0.0410, *SE* = 0.0109, *t*(5886) = −3.77, *p* = 0.0005). ISC-HR at high SNR was also significantly higher than low SNR (Δ^^^ = 0.0287, *SE* = 0.0075, *t*(5886) = 3.83, *p* = 0.0004).

#### Medium SPL (moderately loud environments)

ISC-HR at high SNR was significantly higher than at low or medium SNR (Δ^^^ = 0.02875–0.03086, *SE* = 0.00831–0.00903, *t* = 3.416–3.458, *p* = 0.0016–0.0019), while low and medium SNR did not differ (Δ^^^ = 0.00211, *SE* = 0.00832, *t*(5886) = 0.254, *p* = 1.000).

#### High SPL (loud environments)

ISC-HR values were generally lower across all SNR levels. Only the contrast between high and medium SNR reached significance (Δ^^^ = −0.01818, *SE* = 0.00744, *t*(5886) = −2.443, *p* = 0.0437); other contrasts were not significant (high–low: *p* = 0.2264; low–medium: *p* = 1.000).

This pattern suggests that low-to-medium SPL combined with sufficient acoustic clarity (medium-to-high SNR) supports stronger physiological alignment, whereas excessively loud environments are linked to reduced synchrony. In particular, the noisiest conditions produced ISC-HR levels comparable to those observed during dispersed interactions, indicating that such environments may coincide with less interactive or socially engaging moments.

## 3 Discussion

In this study, we demonstrate that interpersonal physiological synchrony (IPS), as indexed by inter-subject correlation of heart rates (ISC-HR), emerges robustly in everyday social interactions and is modulated by both social and environmental factors. Across three multi-day trips, heart rate synchrony was consistently higher when participants were in close physical proximity, when they interacted with familiar peers, and during activities that fostered shared attention and engagement, such as group games and attending lectures. In contrast, synchrony was reduced during dispersed interactions where opportunities for reciprocal social exchange or shared sensory input were more limited. Moreover, the acoustic environment further modulated ISC-HR: favorable listening conditions (low-to-medium SPL and medium-to-high SNR) enhanced synchrony, whereas noisy conditions reduced it, suggesting that environmental listening conditions influence the degree of physiological alignment, potentially due to increased cognitive effort and attentional load. Importantly, these effects were replicated across the three studies (conducted across different years), and could not be attributed to higher global arousal as we found no association between ISC-HR and mean heart rate. These findings show that IPS is not restricted to high-arousal or controlled laboratory settings, but also emerges spontaneously in naturalistic, everyday settings, making it a potential marker of social engagement under typical, low-intensity social conditions.

IPS is a well-documented phenomenon that occurs when individuals engage in both direct and indirect interaction[5, 26, 41, 42]. However, participant interactions in most of studies investigating IPS have been confined to laboratory settings, potentially limiting the ecological validity of the findings; or high arousal settings, which may drive synchrony through large variations in heart rates[16, 23]. To better understand IPS in naturalistic contexts, we analyzed heart rate (HR) data collected via wearable sensors during three different multi-day real-world trips, capturing both scheduled activities and free-roaming exploration.

Our results demonstrate that people’s heart rates synchronize when they are in close physical proximity. GPS-based measures of spatial distance revealed that both group- and dyad-level ISC-HR were higher when participants were in close physical proximity of each other, compared to time-misaligned control analyses. This pattern was robust across all three trips with different participants, suggesting that shared physical space alone can facilitate physiological coupling in real-world environments. However, the result was less robust across trips for the dyad-level close–far comparison. While dyads in Trips 2 and 3 showed higher ISC-HR when they were close together than when they were far apart, the difference was not significant for Trip 1. Notably, the effect sizes for the dyad-level (Trip 1: *d* = 0.76; Trip 2: *d* = 0.57; Trip 3: *d* = 0.79) comparisons were smaller than those observed at the group level (Trip 1: *d* = 2.94; Trip 2: *d* = 1.9; Trip 3: *d* = 1.35). This may be because close proximity at the dyadic level does not necessarily indicate that partners were interacting or processing the same information, whereas proximity at the group level more likely reflected participation in a shared activity. This weaker effect in Trip 1 may thus be explained by lower effect size and the substantially smaller amount of usable data. Specifically, the number of exact matching pair segments with sufficiently long continuous data for the close versus far conditions was much lower in Trip 1 (1262 vs. 1163) compared with Trips 2 (9542 vs. 8682) and 3 (2904 vs. 3385).

Beyond physical proximity, social familiarity was also found to modulate IPS. Pairs who knew each other prior to the trip exhibited higher ISC-HR than unfamiliar pairs, while controlling for physical proximity, indicating that pre-existing familiarity enhances alignment. This effect may reflect greater shared engagement to common stimuli or enhanced face-to-face interaction among familiar individuals. Interestingly, physiological synchrony was robust enough to capture this effect, despite all participants (familiar and unfamiliar) being in close proximity, and hence exposed to common environmental stimuli.

Examining ISC-HR under different social contexts revealed that people’s heart rates tend to synchronize during close-proximity interactions (e.g., group discussions) and when co-attending to common stimuli (e.g., watching a lecture or performance), but not during dispersed interactions (e.g., sitting scattered in a restaurant). The observed group ISC-HR during stimulus-driven events aligns with previous laboratory research showing heart-rate synchrony between people listening to common narrative stimuli[26] and real-world studies on physiological entrainment to music during live concerts[43]. Our findings thus indicate the context-dependent nature of physiological synchrony, which could be related to differences in participants’ levels of attentional engagement. For instance, during collaborative group activities, participants may have felt more motivated to engage in frequent, dynamic verbal exchanges compared to less interactive settings. This heightened reciprocity in interpersonal communication might have contributed to the observed higher degree of alignment in physiological responses.

Further, we examined how variations in the acoustic environment, characterized by SPL and SNR, were associated with the strength of IPS. The interaction between SPL and SNR indicated that the association between SNR and physiological synchrony depended on the overall loudness of the environment. In quieter settings (low SPL), ISC-HR was highest at medium SNR, while at moderate SPL ISC-HR was highest for high SNR. In louder environments (high SPL), ISC-HR was generally lower across all SNR levels, with values comparable to those observed during dispersed interactions. This may indicate that very loud settings coincide with less interactive activities, or create conditions that make social engagement more challenging. Importantly, because our data are observational, we cannot determine whether these associations reflect a direct effect of the sound environment on physiological synchrony or whether they are driven by the types of activities that typically occur under different acoustic conditions. One limitation is that we did not measure cognitive effort or listening engagement directly, for example via experience sampling. Thus, while our findings show that SPL and SNR are associated with variations in IPS, the underlying mechanisms remain speculative.

The current study thus demonstrates the broader applicability of ISC-HR as a method for quantifying interpersonal synchrony (IPS) beyond controlled laboratory settings or high arousal events. By utilizing wearable sensor data, we have shown the robustness of ISC-HR as a tool for assessing collective, socially and environmentally modulated physiological states. This approach opens new possibilities for studying social dynamics in real-world environments, potentially serving as a marker for monitoring social engagement during real-world interactions.

While the study’s ecological validity is a key strength, it also presents with some inherent limitations. First, there is an inevitable trade-off in data quality when conducting research in real-world settings, as HR data were collected using wristbands instead of the raw cardiac activity data typically obtained from electrocardiograms (ECG) in laboratory environments. The wristbands are less accurate and more susceptible to motion artifacts and variations in activity types. Additionally, optimal GPS tracking using mobile phones requires an unobstructed view of the sky[44], whereas in our case, the participants spent considerable time indoors. These limitations in data quality necessitated extensive pre-processing decisions (cleaning, re-sampling, and interpolation). The scarcity of prior literature addressing these methodological challenges made it particularly difficult to establish standardized methods for handling such data in real-world contexts.

There was also limited control over how and when the participants interacted with each other aside from having the timestamps of planned activities. Consequently, we could not precisely identify the type of interaction each participant was engaged in when in close physical proximity to others. This constraint made it challenging to distinguish between reciprocal interaction and stimulus locking as drivers of IPS, leaving us to make inferences and generalize based on event labels. Future research should specifically examine how different types of interactions — such as conversation, shared attention, or cooperative tasks — affect the strength of ISC-HR, to better understand the mechanisms driving physiological synchrony. Such investigations could help isolate the specific social and cognitive factors that contribute to physiological alignment.

The present findings highlight the potential of IPS as a scalable, ecologically valid index of shared engagement in everyday life. By combining wearable sensing, environmental acoustics, and social context mapping with measures of interpersonal synchrony, our approach provides an index of collective behaviour in real-world environments. Importantly, our results are robust, replicating across three different trips and studies. This framework has broad applicability for quantifying real-world social dynamics. Measures of physiological synchrony can be used to track moment-to-moment fluctuations in collective engagement, to study how social connection emerges and evolves across contexts such as education and collaboration, and to index listening effort and participation in individuals with sensory challenges such as hearing loss. More broadly, combining physiological coupling with environmental and behavioural data offers a pathway toward social sensing - a data-driven approach to characterizing how humans co-regulate their physiological states within complex social and sensory environments.

## 4 Methods

### 4.1 Participants and ethics

Seventy-two participants (60 male; age range 19–32 years, *M* = 23, s.d. = 2.09) took part in three four-day study trips to New York City (23, 24, and 25 participants per trip). Participants were recruited through three editions of the Audio Explorer Challenge[37], a competition hosted by Oticon A/S. They entered the competition in pre-formed teams consisting of 2 to 5 individuals, with the winning teams rewarded with participation in a study trip to New York City. Participants provided informed consent for the use of anonymized data in aggregated analyses and agreed that data could be stored on Oticon A/S-owned secure servers. Additional consent was obtained for access to GPS location and health data via Apple HealthKit. All procedures followed a privacy-by-design framework in compliance with the General Data Protection Regulation (EU 2016/679). The Danish Capital Region Scientific Ethical Committee determined that this study did not constitute health research under the Act on Research Ethics Review of Health Research Projects and therefore did not require ethical approval (reference number F-25048245).

### 4.2 Study design and data collection

Data were collected during three separate four-day trips to New York City. Participants engaged in a combination of pre-planned group activities and unstructured free time, enabling a wide range of real-world social interactions. Continuous multimodal data were collected between 9:00 AM and 10:00 PM each day. Every 20 seconds, hearing aids (Oticon Opn™ S) recorded estimates of the surrounding sound environment, while wristbands (Garmin VivoSmart 4) measured instantaneous heart rate at each detected heartbeat using pulse plethysmography. Smartphones (iPhone 7) recorded GPS location data. Event timelines were documented by the research team and combined with GPS data to label social context and identify periods group interaction based on physical proximity. Acoustic feature extraction and signal-processing details for the hearing aid microphones are provided in Supplementary Information S1. Technical characteristics and limitations of the heart rate sensors are described in Supplementary Information S2.

### 4.3 Pre-Processing

#### 4.3.1 GPS data

Location logs with invalid coordinates were removed. Data were restricted to the Greater New York Area using predefined latitude (40.0–45.0) and longitude (−75.0 to −72.0) boundaries. Spatial distances and time differences between consecutive samples were computed using the Haversine formula to derive speed estimates. Movement status was classified as stationary or moving using a 2 m/s threshold, with stationary points retained for analysis. Large spatial jumps (*>* 30 m within 20 s) were identified and interpolated to reduce GPS artifacts.

#### 4.3.2 Heart rate data

Instantaneous heart rate data were resampled to 10 Hz and interpolated using Piecewise Cubic Hermite Interpolating Polynomials (PCHIP), which minimizes error in wearable heart rate signals[45]. Data were segmented into non-overlapping 5-minute epochs corresponding to events, group periods, and dyadic interaction periods.

The 5-minute window length was selected based on systematic comparisons with 3- and 10-minute windows across all datasets. Shorter windows were unsuitable due to the low sampling rate of physiological and acoustic measures, whereas longer windows reduced sensitivity to short-term interpersonal dynamics. The 5-minute window provided the best balance between temporal resolution, data quality, and participant inclusion, yielding consistent data retention across trips (91.8%, 87.7%, and 92.4%).

##### Analysis

#### 4.3.3 Physical proximity

Physical proximity was assessed at both group and dyadic levels using GPS data. Group periods were defined as intervals of at least 30 minutes during which the average interpersonal distance among participants was below 100 m. This threshold was empirically motivated: during known close-proximity events (e.g., meals, presentations), mean group distances ranged from 0.08–0.15 km, whereas free-time periods showed substantially larger and more variable distances (0.97–2.45 km). The 100 m criterion thus reliably distinguished structured group activities from dispersed periods while accommodating spatial spread within shared environments.

At the dyadic level, participant pairs were classified as close when separated by less than 20 m and far when separated by more than 1 km for periods lasting at least 15 minutes. The 20 m threshold reflects observed proximity during seated or constrained interactions and accounts for the typical horizontal accuracy of smartphone GPS in urban settings (7–13 m)[44].

#### 4.3.4 Social familiarity

Social familiarity was defined based on pre-existing team membership. Participants who entered the study as part of the same team were classified as familiar, whereas participants from different teams were classified as unfamiliar. Familiarity analyses were restricted to periods of close physical proximity. Specifically, we categorized interactions within the group as familiar interactions and those involving members of different teams as unfamiliar interactions. This distinction allowed us to explore the relationship between physical and social proximity and assess how team membership influenced patterns of interaction during group settings.

### 4.4 Quantifying interpersonal heart rate synchrony

Interpersonal physiological synchrony was quantified using inter-subject correlation of heart rate (ISC-HR), a measure used to assess temporal alignment of physiological signals across individuals exposed to shared contexts or stimuli[26, 27]. ISC-HR captures similarity in slow heart rate fluctuations over time and provides a simple, interpretable measure that is robust to noise and well suited for long-duration, naturalistic recordings. A detailed description of signal segmentation, normalization, and ISC-HR computation at the group, dyadic, and familiarity levels is provided in Supplementary Information S3.

Heart rate time series were segmented into non-overlapping 5-minute windows and z-scored within each segment to remove inter-individual differences in mean heart rate and variance. ISC-HR was computed as the Pearson correlation between time-aligned heart rate signals from participant pairs, yielding one degree of synchrony value for each segment.

ISC-HR was computed at multiple analytical levels. At the group level, each participant’s heart rate segment was correlated with those of all other participants present in the same group period, and correlations were averaged to yield a single ISC-HR value per participant. At the dyadic level, ISC-HR was computed for participant pairs during periods of close proximity and during periods when pairs were spatially separated. For social familiarity analyses, ISC-HR was computed separately for familiar and unfamiliar pairs during periods of close proximity.

To assess whether observed ISC-HR exceeded chance levels, we implemented a time-misaligned shuffled control analysis in which heart rate segments were randomly paired across participants such that paired segments did not overlap in time. The number of random pairings matched that of the observed time-aligned data. ISC-HR from time-misaligned pairs provided a control measure of synchrony expected in the absence of shared temporal structure. The full procedure for time-misaligned control analysis is described in Supplementary Information S4.

#### 4.4.1 Modeling the effects of sound environment on ISC-HR

To assess whether the acoustic environment modulated interpersonal heart rate synchrony (ISC-HR), data from all three trips were pooled. For each group period, median sound pressure level (SPL) and signal-to-noise ratio (SNR) were computed across participants for each 5-minute segment[46]. SPL and SNR were categorized into tertiles (low, medium, high) to summarize the acoustic context experienced during group interactions.

Group-level ISC-HR was modeled using linear regression, predicting mean ISC per segment from SPL level, SNR level, and their interaction. In an additional model, mean heart rate (HR) was included as a covariate to account for general arousal. Model comparisons were performed using ANOVA, Akaike Information Criterion (AIC), and changes in *R*^2^. Full model specifications, categorization thresholds, and statistical results are reported in Supplementary Information (S5, Table S1).

### 4.5 Statistical analysis

Statistical analyses tested whether ISC-HR exceeded time-misaligned control levels and whether synchrony varied as a function of physical proximity, social familiarity, social context, and acoustic environment. Assumptions of normality were assessed prior to testing, and appropriate parametric or non-parametric tests were applied. Effect sizes were reported throughout. Additional details regarding statistical testing, model comparisons, and evaluation metrics are provided in Supplementary Information S6.

## Data availability

The dataset underlying this Article is available at https://osf.io/cswt2/overview?viewonly=364e00ef2c4e4b17a34334d43cf5a3d8

## Code availability

The code for replicating the analyses underlying this Article is available at https://osf.io/cswt2/overview?viewonly=364e00ef2c4e4b17a34334d43cf5a3d8

## Acknowledgements

This work is supported by the Carlsberg Foundation’s Semper Ardens: Accelerate grant, “Self, other, and we: mechanisms and dynamics of self-other integration in social observation and interaction”, CF22-1251.

## Author Contribution

H.H., J.H.C., and I.K. designed the study. J.H.C. collected the data. H.H. performed data preprocessing and analysis with input from I.K., J.H.C., J.M.S. I.K. and H.H. wrote the manuscript. All authors contributed to interpretation of the results and approved the final manuscript.

## Ethics approval and consent to participate

Participants submitted their consent agreeing to the use of their anonymized data (i.e., no personal identifiers were available) for research purposes on aggregated levels (i.e. no single-case investigation are performed) and agreed that data could be stored on Oticon A/S-owned secure servers. In addition, participants gave specific consent permitting access to their GPS location and health data from Apple HealthKit. All data collection and storage were conducted in a ‘privacy by design’ manner in accordance with the General Data Protection Regulation (EU regulation 2016/679). The Danish Capital Region Scientific Ethical Committee found that this study was not considered a health research study, according to the Act on Research Ethics Review of Health Research Projects, and therefore did not require ethical approval, reference number: F-25048245 (https://www.nvk.dk/forsker/naar-du-anmelder/hvilke-projekter-skal-jeg-anmelde).

