## Supplementary Information for "Heart rate synchrony as a marker of real-world social engagement: modulated by proximity, social familiarity, and acoustic environment"

This PDF file includes:

- Supplementary Notes S1–S8
- Supplementary Figures S1–S5
- Supplementary Tables S1–S4
- Supplementary references

**S1. Acoustic data acquisition and feature extraction.** The hearing aids logged acoustic data, including SPLs and noise floor (NF) estimates. Unlike laboratory settings, where SNR calculations involve well-defined target signals and noise, SNR in real-world settings relies on estimates from the processed acoustic features sensed by hearing aid microphones. These include environmental sounds, mixed speech, and background noise sources (1, 2).

In this study, the hearing aids used low-pass infinite impulse-response filters with a time constant of 63 milliseconds to estimate SPL and NF. SNR was determined as the difference between the A-weighted (adjusted to mimic human perception of sound as we are not equally sensitive to all frequencies) for SPL and the NF estimate. The SPL represents the immediate sound environment, while the NF represents slower-varying background noise, derived using a 1-second attack and 0.5-second release time for dynamic changes (3). Hearing aid microphones were clipped onto participants' clothing, around the collar area, for consistent data collection.

**S2. Heart rate measurement characteristics.** The Garmin wristbands use data from optical sensors (pulse plethysmographs) to estimate heart rates in beats per minute (BPM). These sensors emit green light through the skin, which is reflected by red blood cells, allowing the wristband to detect changes in blood flow due to heartbeats. The BPM value provides an average heart rate over short intervals (approximately a few seconds), making the readings suitable for long-term tracking, but provide less precise recordings than electrocardiograms (ECG).

ECG, often considered the gold standard in clinical studies, records the electrical activity of the heart by measuring voltage changes on the skin surface, capturing details like P-waves, QRS complexes, and T-waves (4). In contrast, the optical heart rate monitoring of the Garmin wristbands smoothed out small heart rate variations. This method is sensitive to motion artifacts (5) and the participant's physical activity, which can lead to inconsistent RR-intervals not necessarily caused by ectopic beats. As a result, heart rate variability (HRV) analyses were not possible with this dataset. For more details, see Garmin's official heart rate monitoring guide (link) and accuracy disclaimer (link).

**S3. Detailed quantification of interpersonal heart rate synchrony.** Interpersonal physiological synchrony was quantified using inter-subject correlation of heart rate (ISC-HR), which assesses the similarity of temporal fluctuations in physiological signals across individuals (6, 7). This approach captures shared temporal structure without requiring precise event locking and is robust to noise, making it suitable for long-duration naturalistic recordings.

For each participant, instantaneous heart rate time series were segmented into non-overlapping 5-minute windows. Within each window, signals were z-scored to remove inter-individual differences in mean heart rate and variance. ISC-HR was computed as the Pearson correlation between time-aligned heart rate segments.

ISC-HR was computed at multiple analytical levels:

- **Group-level analysis:** Each participant's heart rate segment was correlated with the time-aligned segments of all other participants present during the same group period. Correlations were averaged across segments and partner participants to yield a single ISC-HR value per participant.

- **Dyadic proximity analysis:** ISC-HR was computed for all participant pairs during periods of close physical proximity and during periods when pairs were spatially separated. Correlations were averaged across segments to yield one ISC-HR value per dyad and condition.

- **Social familiarity analysis:** During periods of close proximity, ISC-HR was computed separately for familiar pairs (participants from the same team) and unfamiliar pairs (participants from different teams).

**S4. Time-misaligned shuffled control analysis.** To estimate baseline synchrony expected in the absence of shared temporal structure, we implemented a time-misaligned shuffled control analysis. For each analytical level, heart rate segments were randomly paired across participants such that paired segments did not overlap in time. The number of time-misaligned pairings matched the number of time-aligned pairings used in the observed analyses.

For group-level analyses, each participant's heart rate segments were paired with randomly selected, non-overlapping segments from other participants. For dyadic analyses, time-misaligned segments were generated separately for each participant pair, preserving the number of segments per condition. ISC-HR values computed from these time-misaligned pairings served as a control distribution reflecting synchrony arising from shared signal properties rather than genuine temporal co-fluctuation.

**S5. Modeling the effects of the acoustic environment on ISC-HR.** Associations between the acoustic environment and interpersonal heart rate synchrony (ISC-HR) were examined using pooled data from all three study trips. For each group interaction period, median sound pressure level (SPL) and signal-to-noise ratio (SNR) were computed per 5-minute segment by aggregating values across participants (8). These measures were derived from hearing-aid microphones sampling the acoustic environment every 20 seconds.

To facilitate interpretation and reduce sensitivity to outliers, SPL and SNR were categorized into tertiles reflecting low, medium, and high acoustic conditions. Categorization thresholds were as follows: SPL — low: 38–67.49 dB, medium: 67.66–81.86 dB, high: 81.88–95.54 dB; SNR — low: −2.32–5.65, medium: 5.67–7.17, high: 7.19–17.86.

Group-level ISC-HR was quantified as the mean ISC across participants within each segment. A series of nested linear models were fit to predict mean ISC-HR. Model 1 included additive effects of SPL and SNR. Model 2 additionally included the

SPL × SNR interaction. Model 3 further included mean heart rate (HR) as a covariate to account for general physiological arousal.

$$\text{Model 1: mean\_ISC} \sim \text{SPL\_level} + \text{SNR\_level}$$

$$\text{Model 2: mean\_ISC} \sim \text{SPL\_level} * \text{SNR\_level}$$

$$\text{Model 3: mean\_ISC} \sim \text{SPL\_level} * \text{SNR\_level} + \text{mean\_HR}$$

| Model | Res.df | RSS | Df | F | Pr(>F) |
|---|---|---|---|---|---|
| 1: SPL + SNR | 5891 | 138.13 | – | – | – |
| 2: SPL × SNR | 5887 | 137.00 | 4 | 12.20 | $7.03 \times 10^{-10}$*** |
| 3: SPL × SNR + HR | 5886 | 136.71 | 1 | 12.40 | 0.00043*** |

**Table S1. ANOVA table comparing nested models predicting mean ISC-HR. Including the SPL × SNR interaction (Model 2) significantly improves model fit over additive effects (Model 1). Adding mean HR (Model 3) further improves fit but explains a smaller proportion of variance.**

Model comparison was performed using ANOVA (Table S1) to assess improvements in fit between nested models. Model quality was additionally evaluated using Akaike Information Criterion (AIC) and changes in $R^2$. AIC values were: Model 1 = -5388.47, Model 2 = -5429.06, Model 3 = -5439.47. Marginal $R^2$ values were: Model 1 = 0.0185, Model 2 = 0.0265, Model 3 = 0.0286. Together, these results indicate that the interaction between SPL and SNR accounts for additional variance in ISC-HR, whereas mean HR contributes minimally, suggesting that general arousal does not fully explain the observed effects of the acoustic environment.

**S6. Statistical testing and model evaluation.** To evaluate interpersonal physiological synchrony (IPS) as indexed by inter-subject correlation of heart rate (ISC-HR), several levels of analysis were conducted, and statistical tests were chosen based on data characteristics and experimental design. Across all analyses, normality of differences was assessed using the Shapiro–Wilk test, and appropriate parametric or non-parametric tests were selected accordingly. Effect sizes and multiple comparison corrections are reported where applicable. All statistical analyses were conducted using `R` (version 2024.04.1) and Python (version 3.8).

***Group-level ISC vs. Time-misaligned shuffled controls.*** To test whether ISC-HR exceeded chance levels arising from time-misaligned signals, we compared group-level ISC-HR against time-misaligned shuffled controls. Differences were first tested for normality using the Shapiro–Wilk test. If normally distributed, paired-sample t-tests were used; otherwise, Wilcoxon signed-rank tests were applied. Effect sizes were reported as Cohen's $d$ for t-tests and rank-biserial $r$ for Wilcoxon tests.

***Dyadic ISC across proximity conditions.*** At the dyadic level, ISC-HR was computed for each pair under three physical proximity conditions: close ($< 20$ m), far ($> 1$ km), and time-misaligned shuffled controls. Normality of residuals was assessed using the Shapiro–Wilk test. Repeated-measures ANOVAs were conducted to evaluate the main effect of proximity. When the assumption of sphericity was violated, Greenhouse–Geisser correction was applied. Post hoc pairwise comparisons were conducted between all proximity conditions, with Bonferroni correction for multiple comparisons. Effect sizes are reported as partial $\eta^2$ for ANOVAs and Cohen's $d$ for pairwise contrasts.

***Social familiarity ISC-HR.*** To assess the effect of social familiarity on physiological synchrony, ISC-HR was compared between familiar (participants who knew each other prior to the trip) and unfamiliar pairs, controlling for physical proximity by including only segments where participants were in the same group. Normality of ISC-HR differences was assessed via Shapiro–Wilk tests. When normality assumptions were met, Welch's t-tests were used; otherwise, Wilcoxon rank-sum tests were applied. Effect sizes are reported as Cohen's $d$ for t-tests and rank-biserial $r$ for Wilcoxon tests.

***Event-level ISC-HR analyses.*** To examine ISC-HR across different social contexts, events were categorized as close-proximity interactions, stimulus-locked interactions, and dispersed interactions. For each event, ISC-HR was averaged across all 5-minute segments for all participant pairs. Differences between observed ISC-HR and time-misaligned controls were assessed using paired t-tests if normality was satisfied, or Wilcoxon signed-rank tests otherwise. Effect sizes were reported as Cohen's $d$.

**S7. Controlling for mean heart rate in event-based ISC-HR analyses.** To ensure that differences in ISC-HR across social contexts were not driven by overall heart rate levels, we conducted additional control analyses. For each event, individual HR time series were demeaned prior to ISC-HR computation, and synchrony was compared against time-misaligned shuffled controls.

We further examined whether mean HR was associated with ISC-HR across events. No significant relationship was observed ($r = 0.04$, $p = .45$; Fig. S1), indicating that ISC-HR reflects shared temporal dynamics rather than global arousal differences.

Finally, we assessed mean HR across event types (Fig. S2). Close proximity and dispersed interactions resulted in similarly elevated mean HR, whereas stimulus-locked events showed slightly lower mean HR. This confirms that ISC-HR differences across contexts cannot be explained by overall heart rate differences.

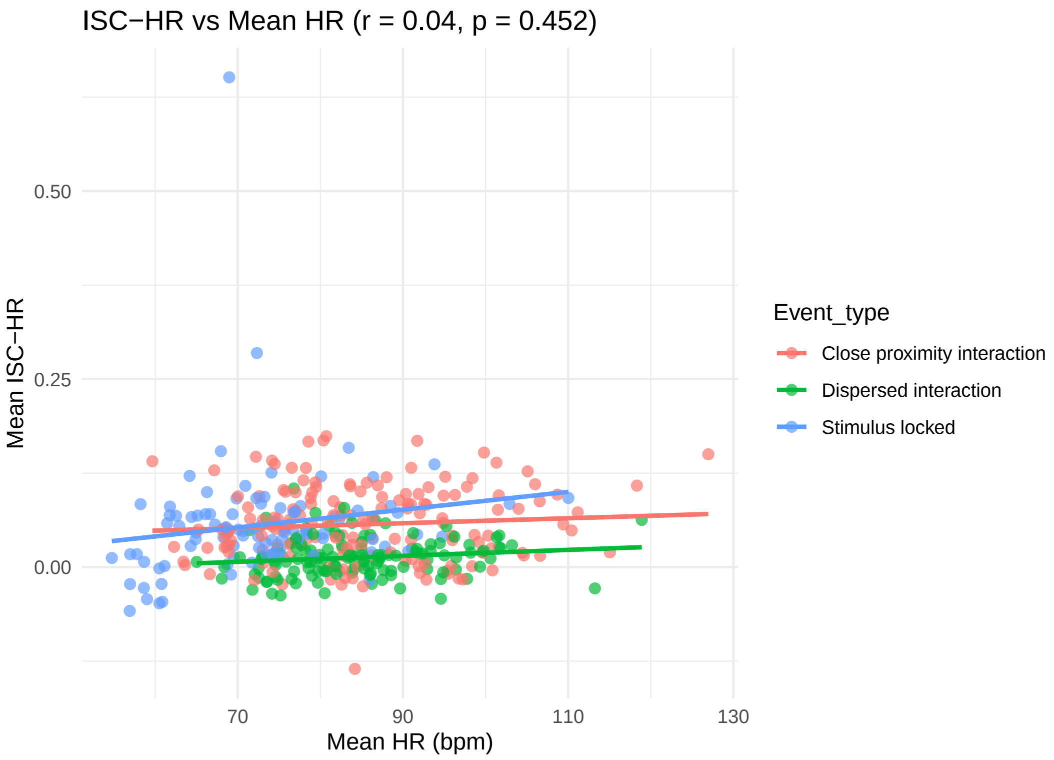


**Fig. S1.** Correlation between mean HR and ISC-HR across events. No significant association was found ($r = 0.04$, $p = .45$).

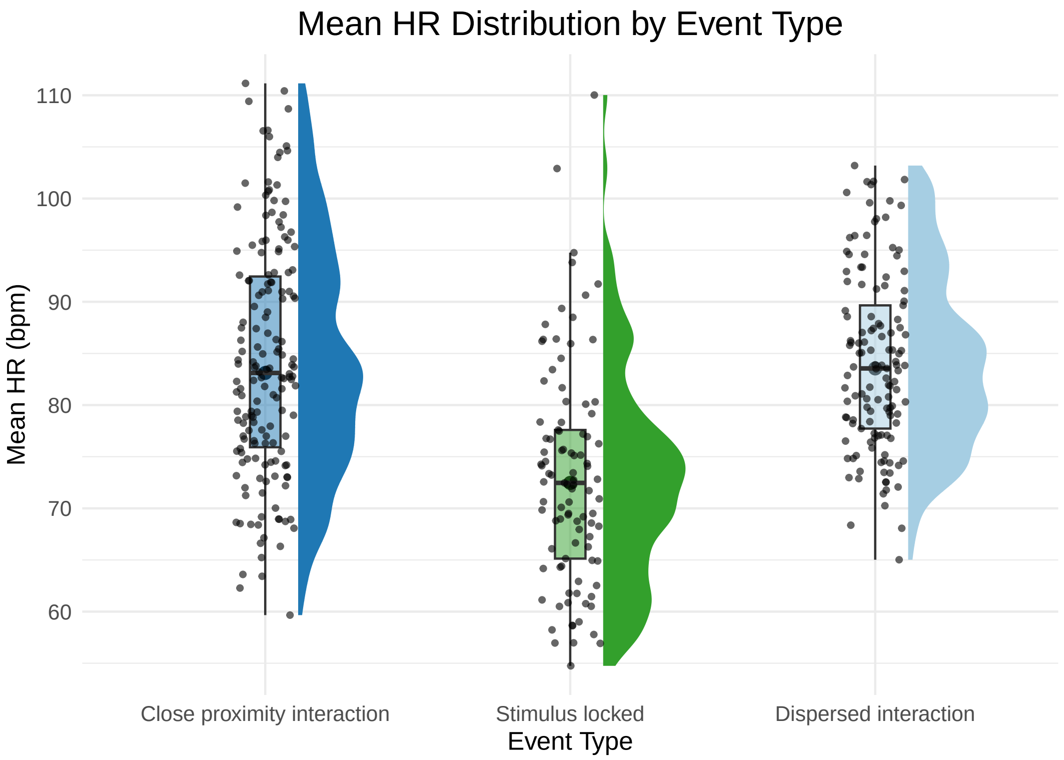


**Fig. S2.** Distribution of mean HR across event types. Close proximity and dispersed events showed similarly high mean HR; stimulus-locked events showed slightly lower mean HR.

**S8. ISC-HR across all events.** For completeness, we report statistical results for each event across the three trips (Tables S2–S4). Raincloud plots comparing original synchrony values with time-misaligned shuffled controls are shown in Figs. S3–S5.

| Interaction Type | Event | Test | Statistic | p-value | Effect size |
|---|---|---|---|---|---|
| Close proximity | Social Dinner 1 | t-test | 2.17 | 0.047 | 0.54 |
| Close proximity | Social Dinner 2 | Wilcoxon | 4.8 | $< 0.001$ | 1.24 |
| Stimulus-locked | Presentation 1 | t-test | 9.2 | $< 0.001$ | 2.0 |
| Stimulus-locked | Presentation 2 | Wilcoxon | 9.0 | $< 0.001$ | 1.01 |
| Dispersed | Casual Reception 1 | t-test | 1.09 | 0.31 | 0.36 |
| Dispersed | Casual Dining 1 | t-test | -0.35 | 0.73 | -0.09 |

**Table S2. ISC-HR statistical results for Trip 1.**

| Interaction Type | Event | Test | Statistic | p-value | Effect size |
|---|---|---|---|---|---|
| Close proximity | Group Game 1 | Wilcoxon | 22 | $< 0.001$ | 1.21 |
| Close proximity | Social Dinner 3 | t-test | 2.43 | 0.028 | 0.61 |
| Stimulus-locked | Show 1 | t-test | 6.06 | $< 0.001$ | 1.35 |
| Stimulus-locked | Presentation 3 | Wilcoxon | 51 | 0.24 | 0.13 |
| Stimulus-locked | Presentation 4 | t-test | 7.51 | $< 0.001$ | 1.64 |
| Dispersed | Casual Dining 2 | t-test | 0.683 | 0.5 | 0.17 |
| Dispersed | Casual Reception 2 | t-test | 1.6 | 0.13 | 0.36 |

**Table S3. ISC-HR statistical results for Trip 2.**

| Interaction Type | Event | Test | Statistic | p-value | Effect size |
|---|---|---|---|---|---|
| Close proximity | Group Game 2 | t-test | 5.51 | $< 0.001$ | 1.26 |
| Close proximity | Social Brunch 1 | t-test | 5.06 | $< 0.001$ | 1.23 |
| Stimulus-locked | Presentation 5 | t-test | 10.92 | $< 0.001$ | 2.33 |
| Stimulus-locked | Presentation 6 | t-test | 6.42 | $< 0.001$ | 1.37 |
| Stimulus-locked | Presentation 7 | t-test | 11.00 | $< 0.001$ | 2.30 |
| Dispersed | Casual Dining 3 | t-test | 2.62 | 0.16 | 0.33 |
| Dispersed | Casual Dining 4 | t-test | 1.23 | 0.27 | 0.24 |
| Dispersed | Casual Reception 3 | t-test | 0.98 | 0.03 | 0.52 |

**Table S4. ISC-HR statistical results for Trip 3.**

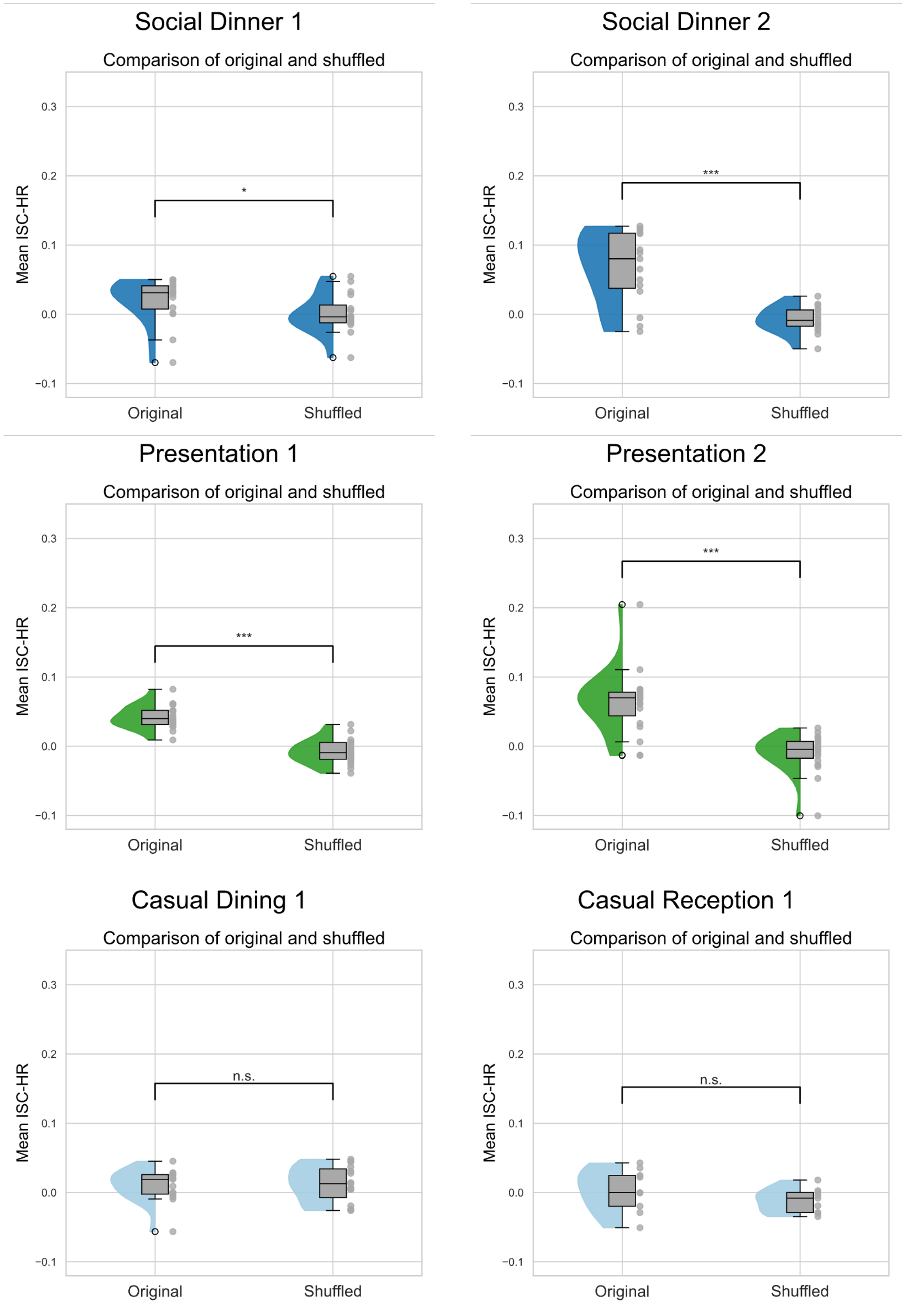


**Fig. S3.** Raincloud plots for Trip 1 showing original vs. time-misaligned shuffled ISC-HR.

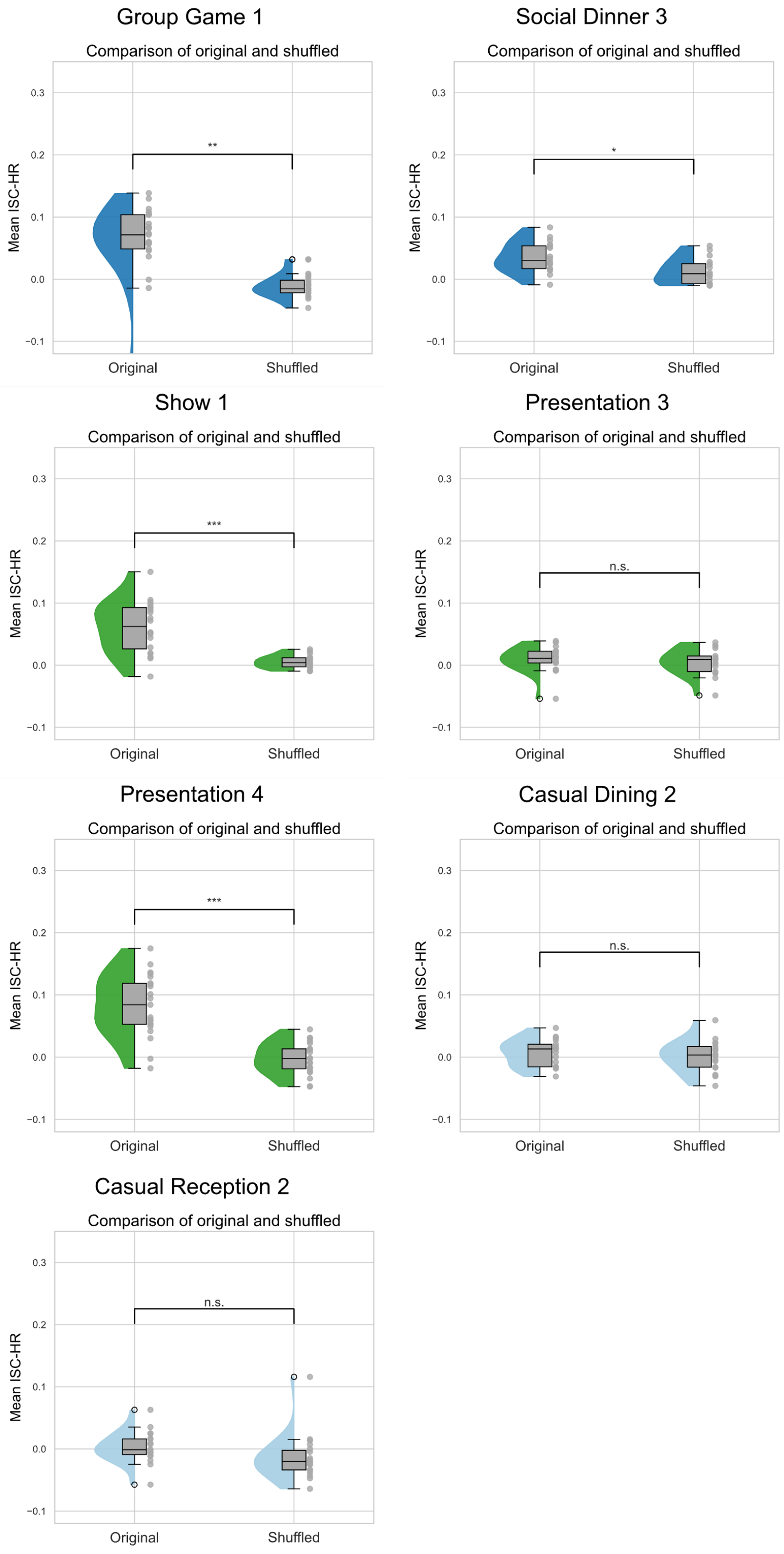


**Fig. S4.** Raincloud plots for Trip 2 showing original vs. time-misaligned shuffled ISC-HR.

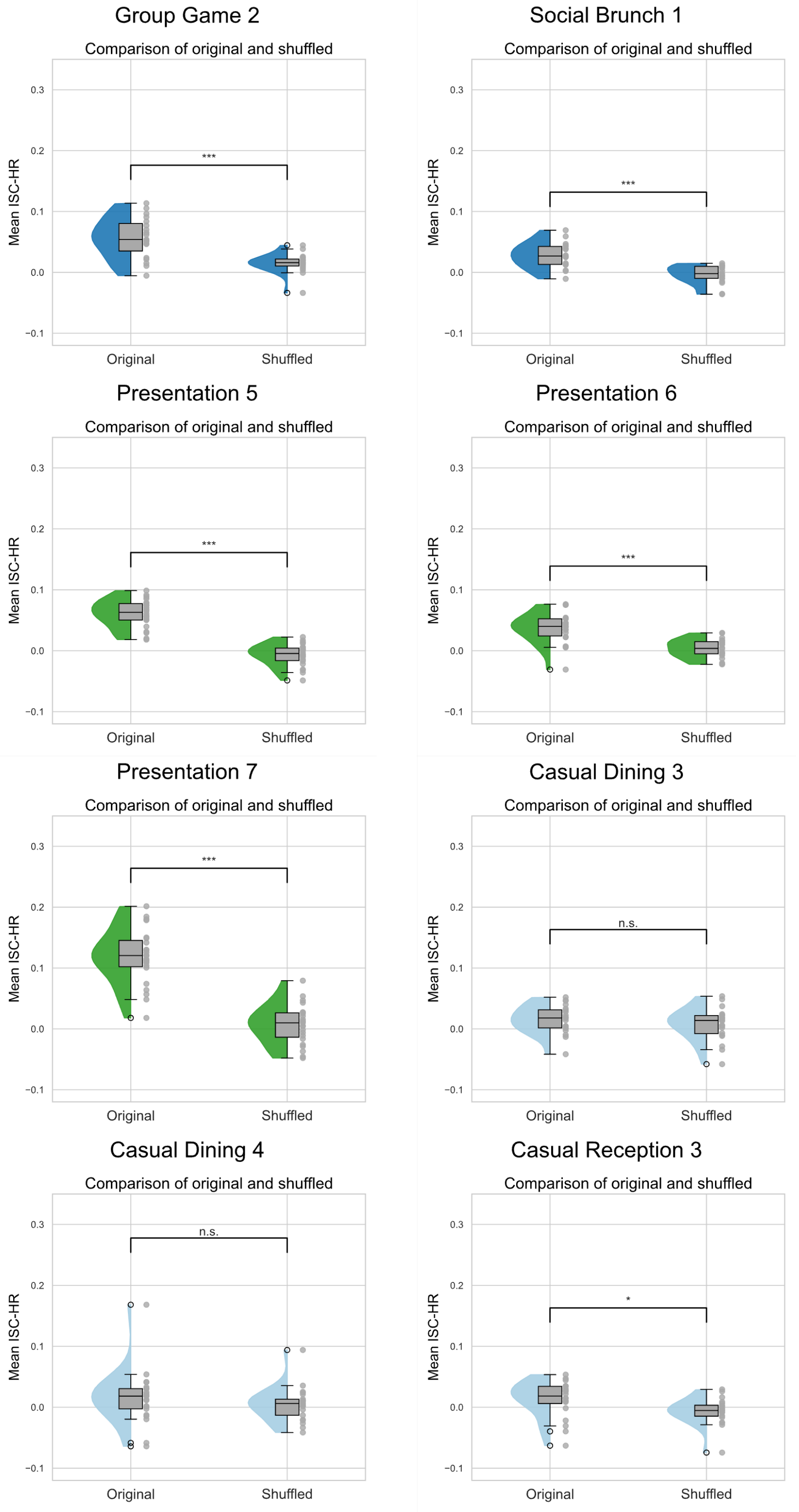


**Fig. S5.** Raincloud plots for Trip 3 showing original vs. time-misaligned shuffled ISC-HR.
